# Effects of early and late life environments on ageing

**DOI:** 10.1101/2021.10.15.464502

**Authors:** Krish Sanghvi, Maider Iglesias-Carrasco, Felix Zajitschek, Loeske E.B. Kruuk, Megan L. Head

## Abstract

Early and late life environments can interact in complex ways to influence the fitness of individuals. Most studies investigating effects of the environment on fitness focus on environments experienced and traits expressed at a single point in an organism’s life. However, environments vary with time, thus the environments organisms experience at different ages may interact to affect how traits change throughout life. Here, we test whether thermal stress experienced during development leads individuals to cope better with thermal stress as adults. We manipulated temperature during both development and adulthood and measured a range of life-history traits, including senescence, in male and female seed beetles, *Callosobruchus maculatus*. We found that favourable developmental conditions increased reproductive performance of females (i.e. silver-spoon effects). In contrast, non-reproductive traits such as lifespan and survival senescence were only affected by adult environments- high adult temperatures decreased longevity and survival. Additionally, developmental and adult environments interacted to affect age-dependent changes in male weight. Overall, our results show that effects of early and late environments can be both sex- and trait- specific, and that a full understanding of how environments interact to affect fitness and ageing requires the integrated study of conditions experienced during different stages of ontogeny.

## Introduction

Early life conditions can act directly on developing phenotypes and in consequence can have both immediate and long-lasting effects on a range of fitness-related traits (Frankenhuis et al, 2019, van de Pol, Bruinzeel et al, 2006). For example, various studies (e.g. Descamps et al, 2008; Klepsatel et al, 2019, 2020; Madsen & Shine, 2000; Muller et al, 2016; Sanghvi et al, 2021; Wong & Kolliker, 2014) have shown that developmental environments can create “silver-spoon effects” (Grafen, 1988; Monaghan, 2008), where individuals who experience a favourable early environment have increased fitness and performance as adults, compared to individuals who experience poor developmental conditions.

The effects of early life environments on fitness are also expected to depend on conditions experienced later in life (Gluckman et al, 2005, Monaghan, 2008). It has been suggested that individuals experiencing certain conditions during development may adjust their phenotype to improve performance when exposed to the same conditions as adults (“environmental matching” or predictive adaptive response). Under this scenario, early life conditions shape the phenotype in response to predicted adult conditions, so that fitness is optimised when environments experienced during development and adulthood match (Bateson et al, 2014; Beaman et al, 2016; Cleal et al, 2007; Hayward & Lummaa, 2013). A special case of the environmental matching hypothesis, the “beneficial acclimation” response, (Huey et al, 1999; Woods & Harrison, 2002), deals explicitly with stressful developmental and adult temperatures. It predicts that stressful temperatures experienced during development acclimate individuals so they perform better when they also experience these stressful temperatures as adults, compared to individuals who only experience stress as adults and not during development (e.g. Bahnrdoff et al, 2016; Deere & Chown, 2006; Kellermann et al, 2017; Kristensen, 2008; Scharf et al, 2015; Scott & Johnston, 2012). Another hypothesis which predicts that adult environments are important for determining later life fitness consequences of early life environments is the “environmental saturation” hypothesis (Engqvist & Reinhold 2016; Pigeon et al, 2019). This hypothesis predicts that in favourable adult environments, all individuals will perform well regardless of their developmental environment, and likewise, that all individuals will perform poorly in bad adult environments. Thus, effects of developmental environments on adult phenotypes are only evident in intermediate adult environments (e.g. Pigeon et al, 2019).

Empirical evidence from studies looking at how early and late life environments interact to affect adult traits is not clear cut, and indicates that the various hypotheses employed to explain the relationships between phenotypic and environmental variation are not mutually exclusive (Pigeon et al, 2019). For instance, some studies find evidence for beneficial environmental matching (Duxbury & Chapman, 2020), while others find other types of interactive (Briga et al, 2017), or only additive (Kleinteich et al, 2015) effects of exposure to poor conditions during both early and late life. Additional complexity to the tangled interactions between early and late life conditions arises from the fact that responses to environmental stimuli can be trait- and sex-dependent (e.g. Duxbury & Chapman, 2020; Helle et al, 2012; Krause et al, 2017; Min et al, 2020; Pigeon et al, 2019; Santos et al, 2021; Scharf, Braf et al, 2015; Stillwell & Fox, 2005). For example, in cichlids reproductive rate is determined only by nutrition during development, while adult growth rate is determined only by nutrition in the adult stage, and clutch size is determined by both developmental and adult life nutrition (Taborsky, 2006). Differences in the way the environment affects different traits may result from energetic and physiological constraints acting on life-history traits (Partridge and Silby 1991), as well as the necessity to allocate resources across traits. Furthermore, differences between males and females in their life-histories and mating strategies mean that selection might favour males and females to respond differently to the same environment (Ceballos & Valenzuela 2011; Maklakov et al, 2009; Stillwell et al. 2010). For example, in seed beetles, males and females respond differently to the presence and density of competitors during the larval stage, leading to sex-specific differences in a variety of life-history traits (Iglesias-Carrasco et al 2020; Sanghvi et al, 2021).

Research shows that, in addition to influencing the absolute expression of traits, the environment can also influence how traits change over an individual’s life, and specifically how they deteriorate with advancing age (i.e. how they senesce) (Balbontin & Moller, 2015; Nussey et al, 2007). Senescence occurs as a consequence of relaxed selection on fitness-related traits in older individuals due to trade-offs between life-history components (Rose & Charlesworth, 2002; Stearns, 1989). However, the rate at which individuals age may depend on a range of factors such as their sex and external environment (e.g. Sanghvi et al, 2021). While there is some support for silver-spoon effects on ageing, with favourable developmental conditions leading to slower reproductive and survival senescence (Hayward, Wilson et al, 2013, Cooper and Kruuk, 2018, Sanghvi et al, 2021), an alternative hypothesis suggest that individuals experiencing good environments may senesce faster due to increased investment in growth and reproduction when young (Adler et al, 2016; Hooper et al 2017; Hunt et al, 2004; Spagopoulou et al, 2020). Additionally, senescence can also depend on the interactions between developmental and adult environments, as seen in studies which test for compensatory growth. Here, organisms which experience poor developmental environments increase their investment in growth in favourable adult environments, although at the cost of increased mortality (Dmitriew & Rowe, 2007; Metcalfe & Monaghan, 2001). While recent research has begun investigating how interactions between developmental and adult environments affect survival and reproductive senescence (Duxbury & Chapman, 2020; Min et al, 2021; Zajitschek et al, 2009), the results do not clearly support one hypothesis (silver-spoon, matching environment, or environmental saturation). Additionally, the age-dependent changes in traits measured are restricted to survival and reproduction, and do not consider other traits which can also change with age and influence fitness.

Here we test whether interactions between heat stress experienced during development and adulthood, if present, are beneficial or not, on various life-history traits, including senescence, in male and female seed beetles (*Callosobruchus maculatus*). Temperature is known to be crucial in determining life-history traits, including senescence, and physiology of ectotherms (Zuo et al, 2012) and is often manipulated in studies which test for silver-spoon effects (e.g. Scharf, Braf et al, 2015), matching environments (e.g. Min et al, 2021), and beneficial acclimation (e.g. Leroi et al, 1994), making it ideal for our experiment. In seed beetles, hot temperatures have been shown to be stressful to survival and reproduction (Fox et al, 2011; Stillwell & Fox, 2005; Stillwell et al, 2007; Vasudeva et al, 2014). However, it is unknown how unfavourable temperatures at different life stages interact with each other to affect senescence, and whether these interactions can be beneficial.

## Methods

### Origin and maintenance of study species

Our stock population of *C. maculatus* was sourced in 2017 from stock kept at the University of Western Australia (see Dougherty et al, 2017 for maintenance details). Once in our lab, stock was maintained for 14-16 generations on cowpea beans (*Vigna unguiculata*) at 24°C - 28°C and 20% - 40% relative humidity. Neither stock nor experimental beetles were provided food or water as adults because they do no need to feed or drink as adults in order to survive and reproduce (Beck & Blumer, 2014).

### Experimental design

To test the effects of developmental and adult temperatures on life-history traits and senescence, we used a split-brood full-sib 2×2 factorial design, in which beetles were assigned to either ‘ancestral’ temperatures (23°C - 25 °C) or ‘hot’ temperatures (33°C - 36°C) during development, and then ancestral or hot temperatures as adults. The ancestral temperature was at the lower end of the temperature range in which the stock had been raised for over 14-16 generations. The hot temperature was a novel and unfavourable environment for this population.

To breed experimental beetles, we collected 86 male and 86 female virgin seed beetles from 150 isolated stock beans, within an hour from when they emerged. Virgin females were randomly paired with virgin males for mating and then given 20-30 beans on which to lay eggs (at 24°C). The beans were checked for eggs every two hours, and those with a single egg laid on them were transferred to individual Eppendorf tubes. If a bean had more than one egg laid on it, the extra eggs were scraped off. Eppendorf tubes containing a bean with an egg, were then randomly assigned to a hot (33°C - 36°C) or ancestral (23°C - 25 °C) developmental temperature for incubation until adults emerged.

On the day of emergence, beetles were weighed (to the nearest 0.01 mg) using a Sartorius Cubis microbalance, and their developmental time (in days) and sex were recorded. Beetles were then assigned to either the hot or ancestral adult temperature treatment, in which they remained until they died. This generated four treatments: *ancestral developmental and ancestral adult (**AA**), ancestral developmental and hot adult (**AH**), hot developmental and ancestral adult (**HA**), and hot developmental and hot adult (**HH**)* temperatures (See Table S1 in Supplement for sample sizes). Males were kept in Eppendorf tubes and weighed every second day. Females were individually mated with a single male from our stock population on the day of their emergence, then transferred to a Petri dish and given 15 new beans each day to lay eggs on. To ensure that all females mated, we observed whether or not the female kicked the male with her hind legs to end copulation (if she did not, we paired her with the same male again after ∼20 minutes). Beans with eggs laid on them were stored in plastic bags, and frozen at -20°C for counting later. Both sexes were checked daily for survival and their adult lifespan was recorded. Details of all traits measured are given in Table 1.

**Table 1.** Summary of traits and models used for analyses. DevT. =Developmental temperature, AdultT. =Adult temperature, Age= Adult age

| Trait | Definition | Model type | Error distribution | Data Transformaton (LMM) / Link Function (GLMM) | Fixed effects | Random Effects |
| --- | --- | --- | --- | --- | --- | --- |
| Emergence success | Likelihood of adults emerging from a bean with an egg laid on it | GLMM | Binomial | Logit | DevT. + Block | Family |
| Development time | Number of days between the laying of an egg, and the emergence of an adult | LMM | Gaussian | Power $((y^\lambda - 1)/\lambda)$<br>Males: $\lambda = -0.133214$<br>Females: $\lambda = 0.2275$ | DevT. + Block | Family |
| Emergence weight | Weight (mg) of beetle on the day of emergence | LMM | Gaussian | Males: None<br>Females: None | DevT. + Block | Family |
| Adult lifespan | Number of days between the emergence of an individual and its death | LMM | Gaussian | Males: None<br>Females: Log<sub>e</sub> (y) | DevT. * AdultT. + Emergence weight + Block | Family |
| Fertility (females) | Likelihood that a female laid at least one egg in her lifetime | GLM | Binomial | Logit | DevT. * AdultT. + Block | None |
| Life time reproductive success (females) | LRS: total number of eggs laid by a female in her lifetime | GLMM | Poisson | Log<sub>e</sub> (y) | DevT. * AdultT. + Adult Lifespan + Block | Family, Observation |
| Age dependent (daily) fecundity (females) | Number of eggs laid by a female each day, measured since day of emergence throughout her lifespan | GLMM | Negative binomial distribution | Log<sub>e</sub> (y) | DevT. * AdultT. * Age (and Age<sup>2</sup>) + Adult Lifespan + Block | Family, Individual ID (and additional models with Age Family and/or Age ID as random effects) |
| Age-dependent<br>weight<br>(males) | Weight (mg) of males measured<br>every alternate day, since day of<br>emergence,<br>throughout his lifespan | LMM | Gaussian | None | DevT. * AdultT. *<br>Age (and Age^2) +<br>Age*Emergence<br>weight +<br>Adult Lifespan +<br>Block | Family,<br>Individual<br>ID |
| Age-dependent<br>survival | Likelihood of adult beetles dying at a<br>given adult age | Cox-Proportion<br>al<br>hazards |  | None | DevT.*<br>AdultT. + Block | Family |

The experiment was conducted over three experimental blocks (Block 1: 26 families; Blocks 2 and 3: 30 families each). The individuals used in the three experimental blocks came from 3 successive generations of stock beetles (i.e. blocks 1, 2, and 3, correspond to generations 14, 15, and 16 respectively). Some experimental beetles escaped, were killed accidentally, or could not have their sex identified accurately during the experiment, and were therefore excluded from analyses of reproduction, weight, and lifespan (57 excluded out of 1381 beetles). The assignment of beetles to the developmental and adult treatments was random except that we tried to assign beetles of each sex from each of the 86 families equally across the four treatments. The observer was blinded to the treatment of beetles during data collection to avoid bias.

### Data analysis

To determine the effects of developmental and adult temperature as well as their interaction on the life-history traits (age-independent traits) and senescence (age-dependent traits) of male and female seed beetles, we used Generalised Linear Mixed-effects Models (GLMM) or Linear Mixed-effects Models (LMM) as appropriate. All analyses were conducted in R v3.5.2 (R Development Core Team, 2011) and models were built using the *lme4* (Bates et al, 2014), and glmmTMB (Magnusson et al, 2017) packages. Model details for each analysed trait are given in Table 1, and in Section B of the online supplement. All models contained experimental block as a three-level fixed effect, and beetle family (i.e. full-sibling groups) as a random effect unless mentioned otherwise. For traits measured prior to the assignment of adult treatments (i.e. emergence success, development time, and emergence weight), we included developmental temperature in the model as a fixed effect. For traits measured after beetles were assigned to adult temperatures (i.e. adult lifespan, female fertility, female lifetime reproductive success, age-dependent (daily) female fecundity, age-dependent male weight, and age-dependent survival), we included both developmental and adult temperatures as well as their interactions as fixed effects. For age-dependent female fecundity and age-dependent male-weight, we included adult lifespan as a fixed effect in the models to account for selective disappearance of beetles. Additionally, for male and female lifespan we included emergence weight as a covariate because larger seed beetles have been previously shown to live longer (Fox et al, 2003). For age-dependent weight of males, emergence weight was also included as a covariate, although here, as an interaction with age, because males at different ages could be affected by their emergence weight in different ways.

We conducted two post-hoc tests to examine how developmental temperatures were affecting age-dependent male weight, within each adult temperature. The first post-hoc model only used data from males who experienced hot adult temperatures, to compare males in AH (ancestral developmental and hot adult) vs HH (hot developmental and hot adult) treatments. The second only used data from males who experienced ancestral adult temperatures, so that males in HA (hot developmental and ancestral adult) and AA (ancestral developmental and ancestral adult) temperatures could be compared.

For female age-dependent (daily) fecundity, we additionally created three models, one which allowed the slopes of different families (of full-sibs) to differ, the other which allowed the slopes of different females to differ, and the third which allowed slopes of both different females and different families to differ. These models were then compared using Akaike information Criteria and log-likelihood ratio tests with the *anova* function in the *stats* package to obtain the best fitting model. This was done to test whether different families and females showed different pattern of reproductive senescence from each other.

For all adult traits, males and females were analysed separately because we were not interested in comparing effects across the sexes (i.e. interactions with sex). For age-independent traits, where models contained interactions, we ran a “full” model with the interactions, as well as a “main-effects” model (if the interactions in the full model were non-significant). The main-effects models excluded non-significant interactions to allow accurate interpretation of the main-effects (Engqvist, 2005). For all linear models, residuals were checked visually to ensure they met assumptions of normality and homoscedasticity. When they did not, the response variable was transformed. To test for overdispersion in our models, we used the function *simulateResiduals* in the package *DHARMa* (Hartig 2020). Where there was evidence for over-dispersion (i.e. for female lifetime reproductive success), we fitted an observation level random effect (Harrison, 2014). To test whether the effects of developmental temperatures were consistent across all blocks, analyses for age-independent traits were followed by post-hoc pairwise comparisons of hot and ancestral developmental temperatures within each block. These comparisons were done using Tukey’s test in the *emmeans* package (Lenth et al, 2019). Effect sizes were calculated as “Hedge’s g” for all two-group comparisons (following equations (1) and (2) in Nakagawa & Cuthill, 2007).

## Results

Our analyses indicated a range of effects of developmental and adult temperature on male and female traits. We describe these results below, and provide a summary in Table 2, with complete model outputs for all analyses presented in the supplementary material (Tables S3 – S18).

**Table 2.** Summary of results for effects of developmental and adult temperatures on all measured traits (ns: P > 0.05). DevT =Developmental temperature, AdultT= Adult temperature, A= Ancestral temperature, H= Hot temperature, Age= Adult age.

| Trait | DevT |  | AdultT |  | DevT* AdultT |  | DevT*Age | AdultT*Age | DevT*AdultT*Age |
| --- | --- | --- | --- | --- | --- | --- | --- | --- | --- |
|  | males | females | males | females | males | females |  |  |  |
| Emergence success (both sexes) | H < A |  |  |  |  |  |  |  |  |
| Development time | H < A | H < A |  |  |  |  |  |  |  |
| Emergence weight | H < A | H < A |  |  |  |  |  |  |  |
| Adult lifespan | H > A | H > A | H < A | H < A | ns | ns |  |  |  |
| Female fertility |  | H < A |  | ns |  | ns |  |  |  |
| Female LRS |  | H < A |  | H < A |  | ns |  |  |  |
| Female age-dependent (daily) fecundity |  | - |  | - |  | - | Yes | Yes | ns |
| Male age-dependent adult weight | - |  | - |  | - |  | Yes | - | Yes |
| Age-dependent survival | ns | ns | Yes | Yes | ns | ns |  |  |  |

### Age-independent traits

Hot temperatures caused lower emergence success, faster development, and lower emergence weight in beetles, than ancestral temperatures. Specifically, hot developmental temperature reduced the *emergence success* of beetles (P< 0.001, Table S3): 676 out of 864 eggs (78%) emerged as adult beetles from the ancestral developmental temperature, while 705 out of 1439 eggs (49%) emerged as adult beetles from the hot developmental temperature. Both males and females that experienced hot developmental temperatures had a shorter *development time* (mean ± SE: males= 23.1 ± 0.1, females= 23.5 ± 0.2 days) than those experiencing ancestral developmental temperatures (mean ± SE: males= 37.8 ± 0.2, females= 37.99 ± 0.2 days) (P< 0.001 for both males and females, Hedge’s g: males= 4.41, females= 4.962, Table S4). Further, both males and females which developed in the hot temperature had a lower *emergence weight* (mean ± SE: males= 3.510 ± 0.030, females= 4.710 ± 0.039 mg) than those developing in the ancestral temperature (mean ± SE: males= 3.750 ± 0.030, females= 5.760 ± 0.042 mg) (P< 0.001 for both sexes, Hedge’s g: males= 0.436, females= 1.4, Table S5).

After controlling for the positive effect of emergence weight (P< 0.001, Section E in supplement) there were contrasting effects of hot temperatures in the developmental versus adult stage on *adult lifespan*. For both males and females, hot developmental temperatures increased adult lifespan (Males: Hedge’s g= 0.10; P= 0.005, Females: Hedge’s g= 0.06; P= 0.036). In contrast, hot adult temperatures decreased male and female adult lifespan (Males: Hedge’s g= 4.820; P< 0.001, Females: Hedge’s g= 2.493; P< 0.001) (Figure 1, Tables S6 and S7). There was no significant interaction between developmental temperature and adult temperature on adult lifespan on either males (P= 0.251), or females (P= 0.531).

**Figure 1.**
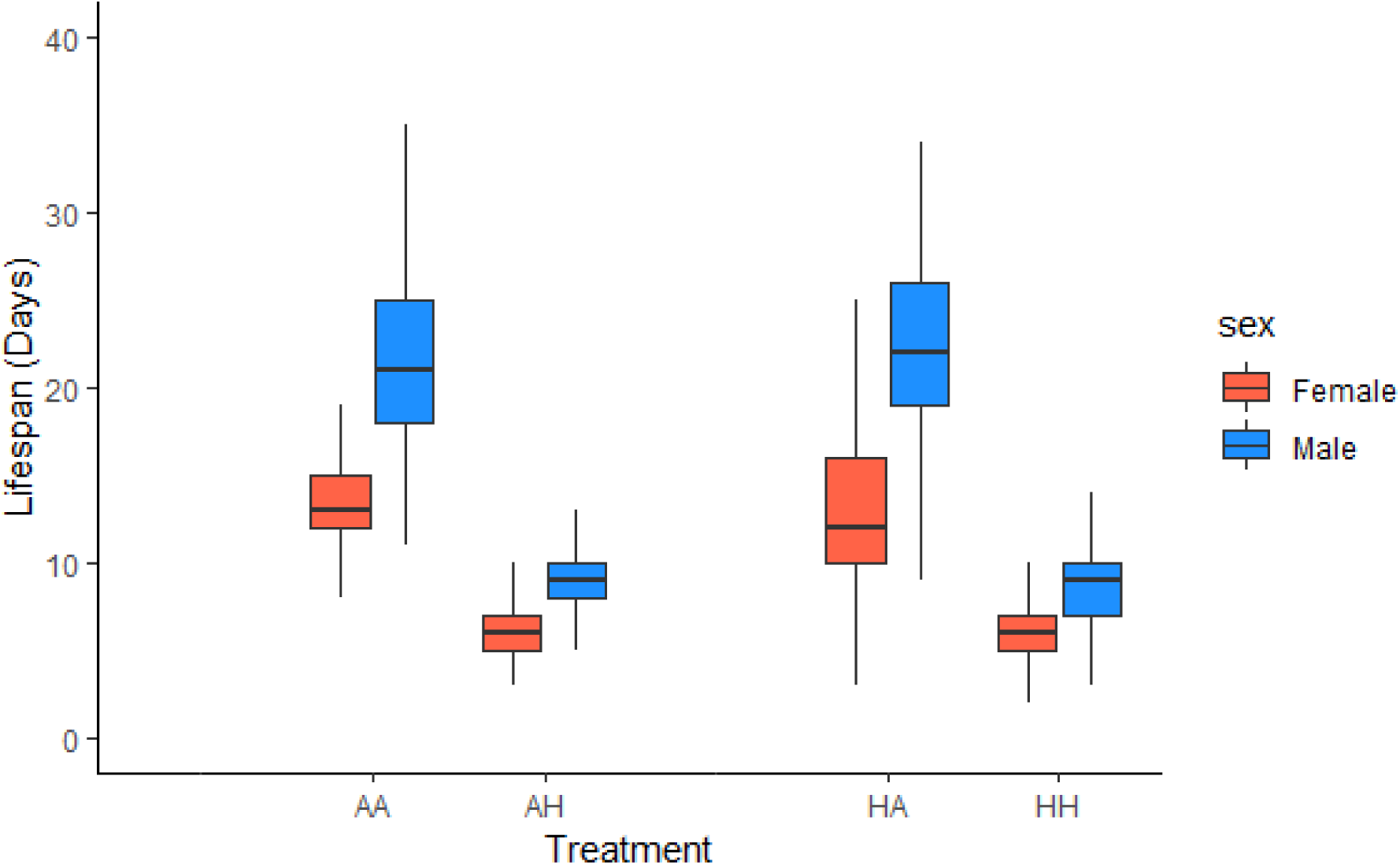
The effects of developmental and adult temperatures on female (red) and male (blue) adult lifespan. *Ancestral developmental and ancestral adult (**AA**), ancestral developmental and hot adult (**AH**), hot developmental and hot adult (**HH**), and hot developmental and ancestral adult (**HA**)* temperatures. Median, interquartile ranges (25,50, 75%) and upper and lower extremes presented.

Females that developed in hot temperatures were less likely to be *fertile* than females which developed in ancestral temperatures (15.5% of females from the hot developmental temperature did not lay any eggs compared to 0.6% of females from the ancestral developmental temperature, P< 0.001, Table S8). Neither adult temperature on its own (P= 0.119), nor the interaction between developmental and adult temperatures (P= 0.986), had a significant effect on female fertility. In contrast, both hot developmental (P< 0.001) and hot adult (P< 0.001) temperatures reduced the *Lifetime Reproductive Success* (*LRS)* of females, compared to ancestral temperatures (Figure 2, Table S9). Although, developmental temperatures (Hedge’s g= 0.945) had a greater effect than adult temperature (Hedge’s g= 0.241) on the LRS of females. There was no effect of the interaction between developmental and adult temperatures on female LRS (P= 0.595).

**Figure 2.**
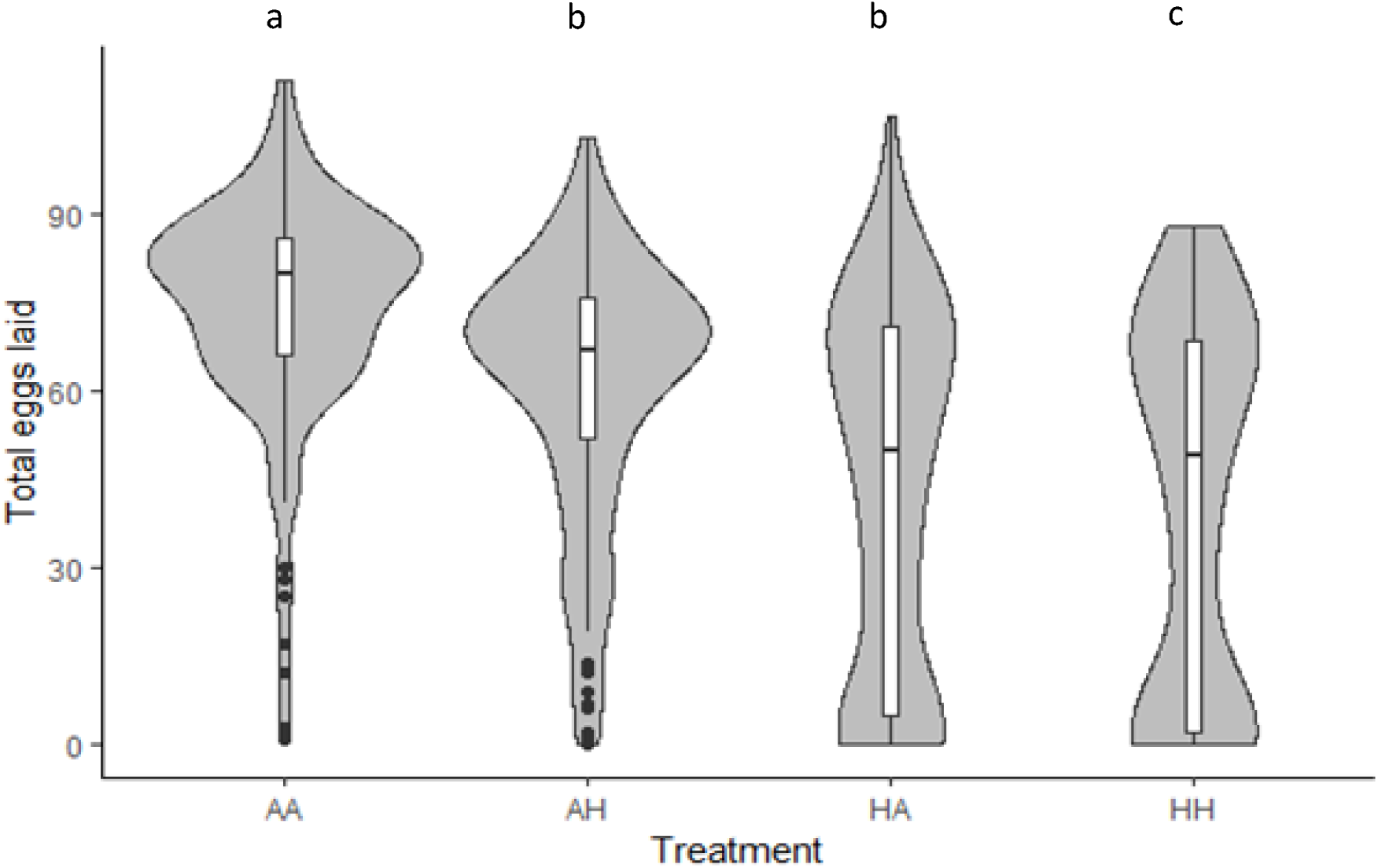
Violin plot comparing the smoothed probability density, median, interquartile ranges (25,50, 75%), and upper and lower extremes presented. of the effects of developmental and adult temperature on female lifetime reproductive success (LRS). *Ancestral developmental and ancestral adult (**AA**), ancestral developmental and hot adult (**AH**), hot developmental and hot adult (**HH**), and hot developmental and ancestral adult (**HA**)* temperatures. Letters ‘a to c’ denote significant differences between pairwise comparisons of groups, with same letters denoting no significant difference.

The effects of developmental environments seen in the results above were consistently found within each block, except for male and female adult lifespan (Table S10).

### Age-dependent traits

Females that experienced either hot developmental or hot adult temperatures showed a steeper decline in *age-dependent (daily) fecundity* as they aged compared to females that experienced ancestral temperatures at either stage (for both stages P< 0.001; Table S11). Females from hot adult temperatures laid a higher number of eggs than females from ancestral adult temperatures at ages one (Hedge’s g= 0.82), two (Hedge’s g= 0.50), and three (Hedge’s g= 0.134) days. After day 3, the effect sizes showed the opposite pattern i.e. females from ancestral adult temperature laid more eggs (Table S12). There was no three-way interaction between developmental temperature, adult temperature, and age (as either linear (P= 0.525) or quadratic (P= 0.226)) to affect the age-dependent (daily) fecundity (Figure 3; Table S11).

**Figure 3.**
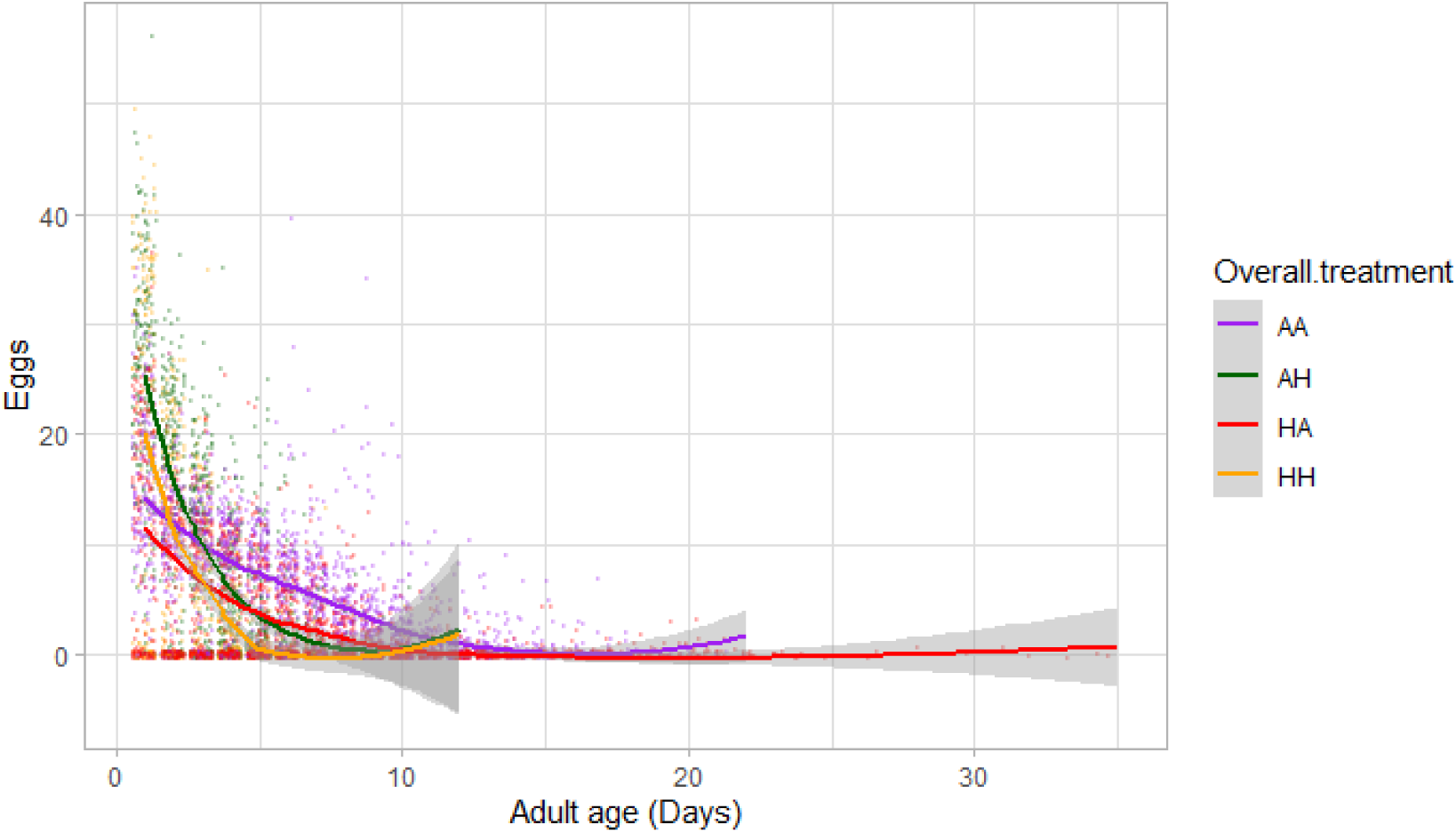
Effects of adult age on female daily fecundity for all four treatments: *Ancestral developmental and ancestral adult (**AA-Purple**), ancestral developmental and hot adult (**AH-Green**), hot developmental and hot adult (**HH-Orange**), and hot developmental and ancestral adult (**HA-Red**)* temperatures. Shaded regions represent 95% confidence intervals.

The model which allowed the slopes for age-dependent (daily) fecundity to vary between families and between females, provided a better fit to the data than models which allowed only the slopes of different families (of full-sibs) to vary, or only the slopes of different individuals to vary, or only the intercepts of families and individuals to vary (Table S13). This suggests both significant between-family and between-individual variation in female reproductive senescence rates.

There was a significant effect of the three-way interaction between developmental and adult temperatures, and age, on age-dependent *male weight* (P< 0.001, Table S14, Figure 4). Post-hoc models, which analysed age-dependent weight data for hot adult and ancestral adult treatments separately (Tables S15, S16), each revealed an interaction between developmental temperature and age. Specifically, in both hot (Table S15) and ancestral (Table S16) adult environments, males who experienced hot temperature during development had a lower rate of decline in weight with age compared to males who experienced ancestral temperatures during development. The difference in average rate of decline in weight per day was greater between HH and AH treatments (0.049 mg per day) than between AA and HA treatments (0.018 mg per day) (Tables S15, S16).

**Figure 4.**
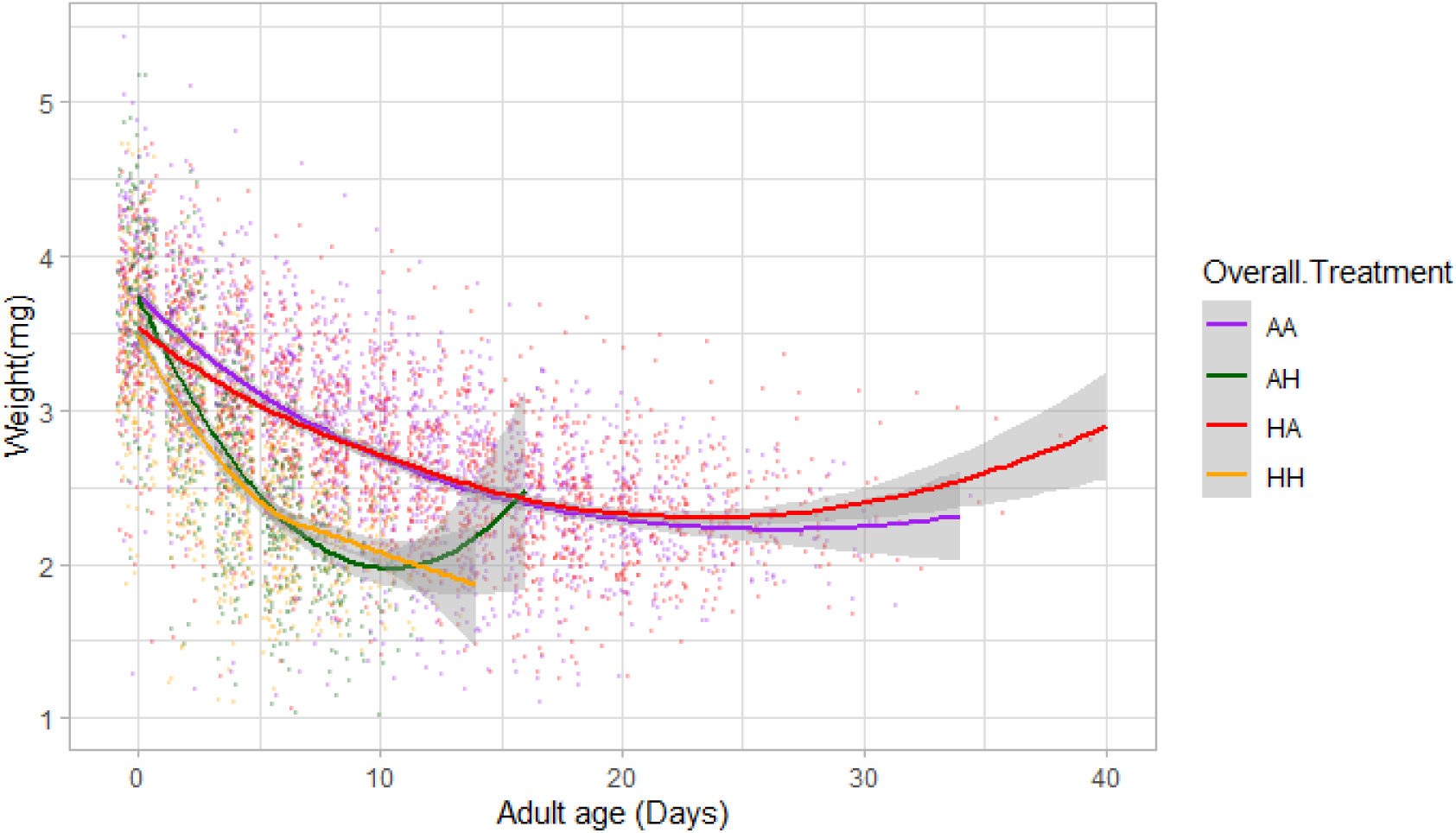
Effects of adult age on weight (mg) in males for all four treatments of: *Ancestral developmental and ancestral adult (**AA-Purple**), ancestral developmental and hot adult (**AH-Green**), hot developmental and hot adult (**HH-Orange**), and hot developmental and ancestral adult (**HA-Red**)* temperatures. Shaded regions represent 95% confidence intervals.

When age-dependent changes in weight were binned by lifespan (Figure S1), they showed that heavier individuals lived longer (See Section E in Supplement). This suggests that the apparent increase in average weight seen in late adult life (evidenced by a significant quadratic effect of age in Table S14 and increase in weight towards the end of life in Figure 4) is due to selective disappearance of lighter beetles (Effect of lifespan: P< 0.001).

For both males (Table S17, Figure 5) and females (Table S18, Figure 6), *age-dependent mortality* (survival senescence) was affected by adult temperature (Males: P< 0.001; Females: P< 0.001), but not developmental temperature (Males: P = 0.210; Females: P= 0.590), nor by an interaction between developmental and adult temperature (Males: P= 0.590; Females: P= 0.820).

**Figure 5.**
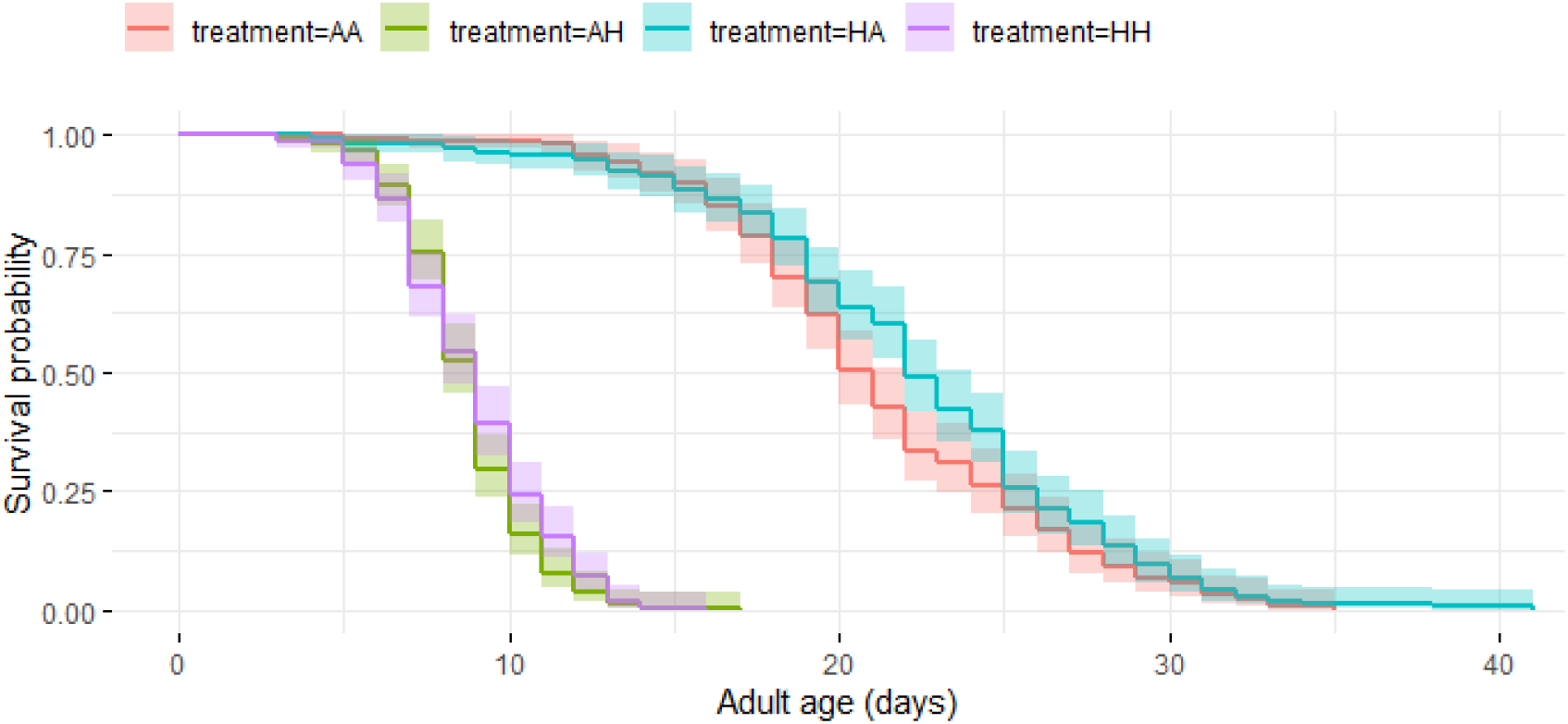
The survival probability of adult males from four treatments with increasing age, namely: *Ancestral developmental and ancestral adult (**AA-red**), ancestral developmental and hot adult (**AH-green**), hot developmental and hot adult (**HH-purple**), and hot developmental and ancestral adult (**HA-blue**)* temperatures, using Kaplan-Meier curves. Shaded regions represent 95% confidence intervals.

**Figure 6.**
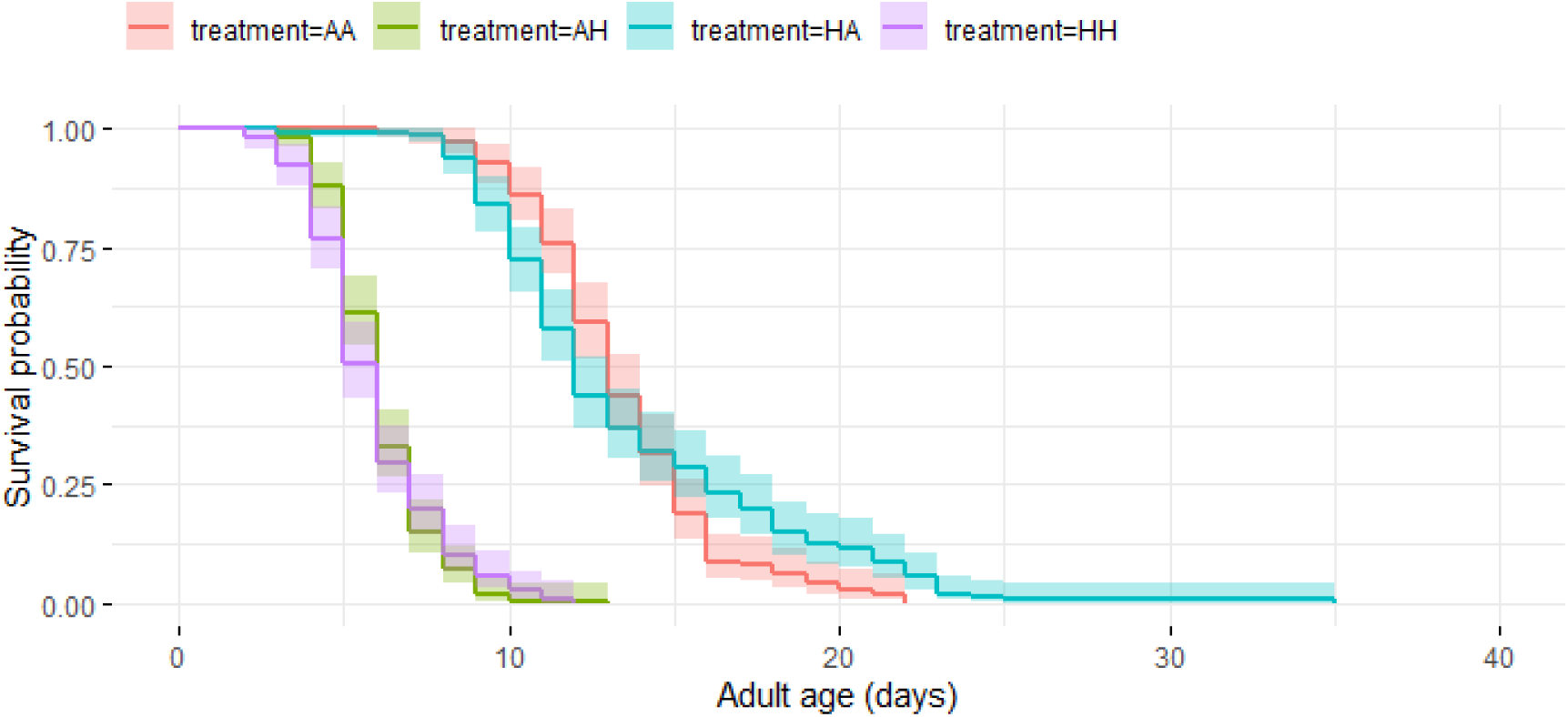
The survival probability of adult females from four treatments with increasing age, namely: *Ancestral developmental and ancestral adult (**AA-red**), ancestral developmental and hot adult (**AH-green**), hot developmental and hot adult (**HH-purple**), and hot developmental and ancestral adult (**HA-blue**)* temperatures, using Kaplan-Meier curves. Shaded regions represent 95% confidence intervals.

## Discussion

Early and late life environments can interact in complex ways to shape adult traits. Consequently, the results of studies testing the effects of exposure to favourable or unfavourable early life conditions both during the development and adulthood are mixed. To improve our understanding of how heat stress experienced at different life stages affects individual phenotypes, as well as to test, whether interactive effects of early and late life environments affect rates of ageing, we subjected juvenile and adult *C. maculatus* to a combination of stressful/hot temperatures and benign/ancestral temperatures. We then measured a range of age-independent and age-dependent traits in both males and females. We found that while female reproductive traits were affected negatively by stressful developmental temperature, somatic traits, such as lifespan and survival senescence, were affected negatively by stressful adult temperature, and weight senescence was affected by an interaction between these environments.

We found that individuals experiencing hot/stressful temperatures during development emerged sooner as adults than those experiencing ancestral temperatures, supporting the metabolic theory of ecology of a faster pace of life (Clarke, 2006). However, accelerated development caused by high temperatures came with costs, such as lower body weight at emergence (which is well known to occur in ectotherms: Zuo et al, 2012), and lower emergence success. Additionally, in females, hot developmental temperatures resulted in reduced reproductive performance (lower LRS and fertility, faster reproductive senescence). These results support the silver-spoon hypothesis for female reproduction, suggesting that independent of the adult environment, favourable developmental environments increase female fitness (see also Cooper & Kruuk, 2018; Sanghvi et al, 2021).

Additionally, when exposed to hot/stressful adult temperatures, females had higher early adult life reproduction (on Days 1 to 3), but shorter lifespans and faster reproductive senescence. These findings support classic life-history theory (Partridge 1987; Stearns, 1989) which proposes a trade-off between early and late life reproduction (Reed et al, 2008), and between survival and reproduction (Kirkwood & Rose, 1999; Marshall et al, 2017). However, our results contrast with a previous study investigating the effects of the developmental density experienced by seed beetles (Sanghvi et al 2021), which showed no effect of high density on the trade-off between female early and late life reproduction, or between female reproduction and survival. This suggests that trade-offs between early and late adult life traits are specific to, and mediated by, the type of environmental parameter being manipulated (e.g. Hammers et al, 2013). In the presence of high temperature, such trade-offs appear to be common across taxa, including seed beetles (Berger et al, 2017; Kim et al, 2020). There are several potential explanations for such patterns. For example, individuals experiencing hot temperatures may adopt a “live-fast die-young” life-history strategy (Robinson et al, 2006). Here, if hot temperatures decrease expected future reproduction and survival, this would be expected to trigger females to terminally invest in early reproduction (Clutton-Brock, 1984), although this increased early life investment would lead to faster senescence in later life (e.g. Gribble et al, 2018). Alternatively, individuals exposed to cooler temperatures might show reduced survival senescence because during early adult life they allocate resources to somatic maintenance rather than reproduction (Kirkwood & Austad, 2000). It is also possible that temperature affects these traits independently, and that the correlations between survival and reproduction are non-causal.

Hot adult temperatures lead to lower adult lifespan and higher age-dependent mortality (survival senescence) in both males and females. The lack of a sex-specific effect on overall survival is not surprising since, again, high temperatures in ectotherms are known to affect longevity as a consequence of increased metabolism (Brown et al, 2004), which is likely to affect males and females in a similar way. More surprising, perhaps, is that the developmental environment did not also affect these somatic traits in a deleterious way. A reason for this could be that selective disappearance of poor quality individuals at the developmental stage of led to an average increase in adult lifespan in hot temperatures, especially in females (See section D in supplement for more information). Another reason may be that somatic traits are only affected by current environments, or that organisms are able to better compensate for poor somatic traits than for reproductive traits, after experiencing unfavourable developmental environments.

In contrast to the other traits, we found that early and late environments interacted to affect age-dependent changes in male weight. This interaction was due to the differences in age-dependent decline in weight between hot and ancestral developmental temperature being greater when adult temperature are hot compared to when they are ancestral (Table S15, S16). Additionally, males who experienced hot temperature at both stages, showed a slower rate of age-dependent loss in weight (Table S15) compared to males who experienced favourable developmental but hot adult temperatures. This result is consistent with the beneficial acclimation hypothesis (Wilson & Franklin, 2002), which is a form of adaptive plasticity. Although, because the absolute differences in the rate of weight loss between males in these treatments of our study is low (only 0.049 mg per day), whether this result is biologically meaningful, remains unknown. Evidence for beneficial acclimation due to an interaction between adult and developmental environments (West-Eberhard, 2003) has been commonly found in species where temperature is manipulated (Leroi et al, 1994; Geister & Fischer, 2007; Reviewed in Wilson & Franklin, 2002). However, previous studies looking at age-dependent traits have not found any evidence for beneficial acclimation effects when manipulating foraging environments (Briga et al, 2019), or diet and temperature (Min et al, 2021). A testable hypothesis for why we found some evidence for beneficial acclimation could be that the allocation of resources towards somatic (i.e. body weight) maintenance, by males who experience developmental stress, occurs at the expense of investment in reproduction.

A recent study (Duxbury & Chapman, 2020), which aimed to test between the silver-spoon and environmental matching hypotheses, manipulated developmental and adult nutrition, and found that female reproductive senescence was affected by an interaction between developmental and adult diets. However, another study that manipulated both adult and developmental temperatures in *Drosophila* found results similar to ours (Min et al, 2020). Specifically, they found that adult temperature had a greater effect on age-dependent survival than developmental temperature, and that hot adult temperatures were deleterious for adult survival senescence. However, they also found an interaction of adult and developmental temperatures to affect age-dependent fecundity deleteriously, which we did not. One reason why we did not find such interactions could be due to the kind of environmental variable that was manipulated (temperature in our study, diet in Duxbury & Chapman, 2020) and different biology of the species studied.

Finally, we also found two population-level patterns in senescence which could have important ecological and evolutionary consequences. First, there were significant differences in rates of reproductive senescence between females, as well as between families (of full-sibs). This was evidenced by models which allowed the slopes of both, different females and families to vary, providing a better fit to the data than models which did not. Individual-level variation in senescence has been shown previously in wild animals (Bouwhuis et al, 2010), and is essential for traits (in our case, reproductive senescence) to evolve via natural selection. Further, the differences in rates of reproductive senescence between families suggest this trait is heritable and thus able to evolve. Second, we found evidence for selective disappearance of lighter males with increasing age. This may have masked individual-level decreases in weight due to heavier individuals surviving longer. Such evidence of individual-level patterns of senescence being masked by population patterns of ageing due to selective disappearance of individuals has previously only been shown in vertebrates (Bouwhuis et al, 2009; Hayward et al, 2013).

## Conclusion

Ours is one of the first studies to test how heat stress experienced at different life-stages interacts to affect individual life-histories and senescence. We show that depending on the trait and the sex measured, either developmental, adult, or the interaction between both environments can affect the resulting phenotype. This suggests that the way environments affect an individual’s phenotypic responses is complex and shows that each trait can follow different trajectories, providing support to different aspects and hypotheses of life-history theory. In consequence, we highlight the importance of measuring fitness-related traits throughout an organism’s life, as well as measuring both physiological and life-history components of fitness in order to understand the holistic effects of environments on individuals. Considering that the adult environment might have stronger influence than developmental environments on traits which are not direct measures of reproduction, we suggest that studies should integrate the effects of early and late life environments to avoid biased results, as well as measure a diverse range of life-history and physiological traits. If traits change differently over time in different environments, it is crucial that studies measure these traits throughout the lifetime of organisms.

