## Supplementary information for "Effects of early and late life environments on ageing"

**Supplementary material**

**Contents**

A. Final sample sizes used to analyse adult traits (page 2)
B. Details of models used to analyse each trait (page 3)
C. Model outputs (page 5)
D. Note on selective disappearance of beetles during emergence (page 20)
E. Note on selective disappearance of lighter males during adulthood (page 21)

Data related to this study can be found on Open Science Framework: https://osf.io/dt5ah/

A. **Final sample sizes used to analyse adult traits**

**Table S1.** Final sample sizes of emerged individuals used to analyse adult traits. *Ancestral Developmental and Ancestral Adult (****AA****), Ancestral Developmental and Hot Adult (****AH****), Hot Developmental and Hot Adult (****HH****), and Hot Developmental and Ancestral Adult (****HA****)* temperatures.

| **Block** | **Treatment** | **Female (N)** | **Male (N)** |
| --- | --- | --- | --- |
| **1**  (Total of 260 eggs setup in Hot Developmental temp. and 260 in Ancestral Developmental temp. from 26 families) | AA | 46 | 45 |
|  | AH | 53 | 42 |
|  | HH | 26 | 21 |
|  | HA | 25 | 21 |
| **2**  (Total of 600 eggs setup in Hot Developmental temp. and 300 in Ancestral Developmental temp. from 30 families)* | AA | 62 | 55 |
|  | AH | 68 | 70 |
|  | HH | 101 | 120 |
|  | HA | 107 | 114 |
| **3**  (Total of 600 eggs setup in Hot Developmental temp. and 300 in Ancestral Developmental temp. from 30 families)* | AA | 40 | 60 |
|  | AH | 50 | 68 |
|  | HH | 29 | 38 |
|  | HA | 28 | 35 |

*More beetles were set up in blocks 2 and 3 because it became apparent after running Block 1 that beetles had higher mortality in the Hot Developmental treatment than in the Ancestral developmental temperature (See Section D in Supplement).

B. **Details of models used to analyse each trait**

***Age-independent traits***

*Emergence success:* We ran a GLMM with a binomial error distribution and “logit” link function. Our binomial response variable was whether a beetle emerged from an egg (“1”) or not (“0”). We included developmental temperature and block as fixed effects.

*Development time:* We created separate models for males and females. For both, a LMM was used in which we included developmental temperature and block as fixed effects and development time as our response variable. Data for both males and females were power transformed using the *car* package.

*Emergence weight:* We analysed males and females separately using an LMM in which we included developmental temperature as a fixed effect, with emergence weight as our response variable.

*Adult lifespan:* We modelled developmental temperature, adult temperature and their two-way interactions as fixed effects using an LMM, separately for males and females. We also included emergence weight of beetles as a covariate in the models to ensure that the effects of temperature on adult lifespan did not result because of differences in emergence weight of beetles. The model which included emergence weight was a better fit than the one which excluded emergence weight for both males (ΔAIC=84) and females (ΔAIC=37). The data for adult lifespan was natural log transformed for females.

*Fertility (Females only):* To analyse whether developmental and adult temperatures affected the likelihood that a female laid at least one egg in her lifetime (henceforth called fertility), we modelled developmental and adult temperature as well as their two-way interaction as a fixed effect, and our binomial response variable was whether a female laid at least one egg (“1”) or not (“0”) during her lifetime. We used the function *glm* in the package *stats* to model the data with a binomial error distribution and “logit” link function. The random effect of family explained no variation, thus was dropped from the model.

*Lifetime reproductive success (LRS):* We modelled the total eggs laid by females across her lifespan, as the response variable using a GLMM. Developmental and adult temperature as well as their two-way interaction were included as fixed effects. We added female adult lifespan as a covariate to account for selective disappearance (van de Pol & Verhulst, 2006). To correct for over-dispersion and non-convergence in this model, the function *glmerControl*(optimizer=”bobyqa”) in the package *lme4* was used to model observational-level random effects.

***Age-dependent traits***

We analysed three age-dependent traits using daily observations of adults following emergence. Because we wished to test how developmental and adult environments interacted with each other, and their effect on ageing rates, these models included three-way interactions between the two temperature treatments and age.

*Female age-dependent (daily) fecundit*y: To check whether there were any effects of developmental and adult temperatures on age-dependent number of eggs laid by females, we fitted a GLMM using the function *glmmTMB* (family=nbinom2) in the package *glmmTMB*, to account for zero-inflation of data (henceforth called Model 1). Eggs laid per day was the response variable. We included two separate three-way interaction of developmental temperature, adult temperature, and age (linear and quadratic) as fixed effects. The model with both linear and quadratic term for age was a better fitting model than a model with just a linear term for age (ΔAIC= 96). We included a random effect of beetle ID to account for repeated measurements made on the same female, and adult lifespan as a fixed effect to account for selective disappearance.

To test whether there were between individual-level differences in age-dependent fecundity between different females, we allowed the slope of each individual to vary independently with age, for number of eggs laid per day, using “Age | Individual I.D.” as an additional random effect in a new model, in addition to all the terms in ‘Model 1’. Further, to measure whether there was between family variation in changes with age in the number of eggs laid, we allowed the slope of each family to vary independently with age for number of eggs laid per day, adding “Age | Family” as an additional random effect in a new model, addition to all the terms in ‘Model 1’ (described above). To further test whether the slopes varied between families after accounting for the effects of between individual variation, we created a model which allowed both between-individual and between-family age-dependent slopes to differ. We then compared each of these three models with the next best fitting model, using a log-likelihood ratio test under a chi squared distribution.

*Age-dependent male weight:* To analyse the effects of developmental and adult temperatures on age-dependent weight of males, we used an LMM. In this “full” model, weight of adult males measured every second day, from emergence until death, was the response variable. In the model, we included two separate three-way interactions of developmental temperature, adult temperature, and age (one term for linear and one for quadratic effects of age) as fixed effects. There was a possibility that the age-dependent changes in weight of males depended on their emergence weight, such that males with lower emergence weights would show a shallower decline in weight. To account for this, we added a fixed effect of an interaction between male age and emergence weight in the model. The model with an interaction term between emergence weight and age was better fitting than the model without this term (ΔAIC= 2483, ΔDF= 2). Additionally, the model with a quadratic term for age was a better fitting model than a model which did not include quadratic effects of age (ΔAIC= 2442, ΔDF= 4). We included a random effect of beetle ID to account for repeated measurements of weight on the same beetles and a covariate of adult lifespan to account for selective disappearance of males. Because the three-way interaction between developmental and adult temperature and age was significant in this model, we conducted two post-hoc analyses. The first post-hoc model only used data from males who experienced hot adult temperatures, to compare males in AH (ancestral developmental and hot adult) vs HH (hot developmental and hot adult) treatments. The second only used data from males who experienced ancestral adult temperatures, so that males in HA (hot developmental and ancestral adult) and AA (ancestral developmental and ancestral adult) temperatures could be compared. All terms in these models were the same as the “full” model described above, except for the term for adult temperature treatment, which was excluded.

*Age-dependent mortality:* To test for the interactive effects of developmental and adult temperature on age-dependent mortality, we analysed males and females separately. We constructed a Cox mixed-effects proportional-hazards model using the function and package *coxme* (Therneau, 2014). This included developmental and adult temperature treatment and their interactions as fixed effects and the adult lifespan of beetles as the response variable.

C. **Model Outputs**

**(**Note: for all tables - predictors of interest shaded in grey, significant P values highlighted in bold.). DevT= Developmental temperature, AdultT= Adult temperature, Age= Adult age, N= number of beetles included in the analysis.

**Table S3.** Effect of Developmental temperature on **emergence success**. Modelled using a GLMM logistical regression with binomial error distribution and “logit” link function. (N = 2303)

| **Fixed effect** | **Estimate** | **SE** | **z** | **P** |
| --- | --- | --- | --- | --- |
| (Intercept) | 1.092 | 0.141 | 7.750 | 0.000 |
| DevT (Hot) | -1.657 | 0.112 | -14.853 | **<0.001** |
| Block2 | 1.731 | 0.177 | 9.754 | <0.001 |
| Block3 | -0.435 | 0.166 | -2.619 | 0.009 |
| **Random effect** | **Variance** | **SD** |  |  |
| Family | 0.173 | 0.416 |  |  |

**Table S4.** The effect of developmental temperature on **development time** for A.) Males (N=703) and B.) Females (N=667). Modelled using LMMs. Power ((y^λ^ -1)/λ)) transformation of data for males (λ= -0.133214) and females (λ= 0.2275)

| **A. Males** |  |  |  |  |  |
| --- | --- | --- | --- | --- | --- |
| **Fixed effects** | **Estimate** | **SE** | **DF** | **t** | **P** |
| (Intercept) | 2.883 | 0.006 | 141.444 | 459.058 | <0.001 |
| DevT (Hot) | -0.299 | 0.004 | 681.140 | -66.730 | **<0.001** |
| Block2 | -0.041 | 0.008 | 100.218 | -5.326 | <0.001 |
| Block3 | 0.026 | 0.008 | 113.269 | 3.255 | 0.002 |
| **Random effects** | **Variance** | **SD** |  |  |  |
| Family | 0.000 | 0.018 |  |  |  |
| **B. Females** |  |  |  |  |  |
| **Fixed effects** | **Estimate** | **SE** | **DF** | **t** | **P** |
| (Intercept) | 5.702 | 0.020 | 118.290 | 291.991 | <0.001 |
| DevT (Hot) | -1.000 | 0.015 | 634.280 | -67.631 | **<0.001** |
| Block2 | -0.145 | 0.024 | 81.267 | -6.045 | <0.001 |
| Block3 | 0.051 | 0.026 | 110.088 | 1.917 | 0.058 |
| **Random effect** | **Variance** | **SD** |  |  |  |
| Family | 0.003 | 0.057 |  |  |  |

**Table S5**: The effect of developmental temperature on **emergence weight** for both A.) Males (N=701) and B.) Females (N=668). Modelled using LMMs.

| **A. Males** |  |  |  |  |  |
| --- | --- | --- | --- | --- | --- |
| **Fixed effects** | **Estimate** | **SE** | **DF** | **t** | **P** |
| (Intercept) | 3.414 | 0.063 | 113.153 | 54.613 | <0.001 |
| DevT (Hot) | -0.269 | 0.040 | 665.854 | -6.773 | **<0.001** |
| Block2 | 0.389 | 0.078 | 84.73 | 4.993 | <0.001 |
| Block3 | 0.494 | 0.081 | 93.533 | 6.123 | <0.001 |
| **Random effect** | **Variance** | **SD** |  |  |  |
| Family | 0.005 | 0.068 |  |  |  |
| **B. Females** |  |  |  |  |  |
| **Fixed effects** | **Estimate** | **SE** | **DF** | **t** | **P** |
| (Intercept) | 5.368 | 0.077 | 115.600 | 69.931 | <0.001 |
| DevT (Hot) | -1.088 | 0.055 | 630.787 | -19.885 | **<0.001** |
| Block2 | 0.488 | 0.095 | 82.582 | 5.112 | <0.001 |
| Block3 | 0.688 | 0.104 | 108.616 | 6.622 | <0.001 |
| **Random effect** | **Variance** | **SD** |  |  |  |
| Family | 0.062 | 0.250 |  |  |  |

**Table S6:** The effect of developmental temperature and adult temperature on **male adult lifespan** for (N=689), after accounting for the effects of emergence weight. Modelled using LMMs. “Full model” shows the parameter estimates and significance values for the model with two-way interactions (highlighted in grey), while the main-effects model shows parameter estimates and significance values for interpretation of only the main-effects (highlighted in grey).

| **Full model** | **Fixed effects** | **Estimate** | **SE** | **DF** | **t** | **P** |
| --- | --- | --- | --- | --- | --- | --- |
|  | (Intercept) | 11.080 | 1.076 | 600.304 | 10.298 | <0.001 |
|  | DevT (Hot) | 1.188 | 0.419 | 641.835 | 2.836 | 0.005 |
|  | AdultT(Hot) | -12.639 | 0.393 | 618.375 | -32.199 | <0.001 |
|  | Emergence weight | 2.433 | 0.281 | 680.817 | 8.673 | <0.001 |
|  | Block2 | 1.572 | 0.563 | 97.871 | 2.791 | <0.001 |
|  | Block3 | 1.366 | 0.590 | 112.129 | 2.316 | 0.022 |
|  | DevT(Hot)*AdultT(Hot) | -0.632 | 0.551 | 617.303 | -1.148 | 0.251 |
|  | **Random effect** | **Variance** | **SD** |  |  |  |
|  | Family | 2.045 | 1.430 |  |  |  |
| **Main effects  model** | **Fixed effects** | **Estimate** | **SE** | **DF** | **t** | **P** |
|  | (Intercept) | 11.222 | 1.069 | 595.990 | 10.498 | <0.001 |
|  | DevT (Hot) | 0.857 | 0.304 | 660.224 | 2.817 | **0.005** |
|  | AdultT(Hot) | -12.960 | 0.275 | 616.865 | -47.055 | **<0.001** |
|  | Emergence weight | 2.439 | 0.281 | 681.753 | 8.692 | <0.001 |
|  | Block2 | 1.586 | 0.563 | 97.992 | 2.818 | 0.006 |
|  | Block3 | 1.375 | 0.589 | 112.349 | 2.332 | 0.021 |
|  | **Random effect** | **Variance** | **SD** |  |  |  |
|  | Family | 2.045 | 1.430 |  |  |  |

**Table S7**. The effect of developmental temperature and adult temperature on **female adult lifespan** (N=635), after accounting for the effects of emergence weight. Modelled using LMMs. “Full model” shows the parameter estimates and significance values for the model with two-way interactions (highlighted in grey), while the main-effects model shows parameter estimates and significance values for interpretation of only the main-effects (highlighted in grey). Log_e (y) data transformation applied.

| **Full model** | **Fixed effects** | **Estimate** | **SE** | **DF** | **t** | **P** |
| --- | --- | --- | --- | --- | --- | --- |
|  | (Intercept) | 2.056 | 0.089 | 547.045 | 23.085 | <0.001 |
|  | DevT (Hot) | 0.073 | 0.036 | 614.761 | 2.037 | 0.042 |
|  | AdultT(Hot) | -0.801 | 0.031 | 576.232 | -26.245 | <0.001 |
|  | Emergence weight | 0.083 | 0.016 | 592.700 | 5.293 | <0.001 |
|  | Block2 | 0.005 | 0.032 | 80.887 | 0.170 | 0.865 |
|  | Block3 | 0.113 | 0.036 | 124.056 | 3.121 | 0.002 |
|  | DevT(Hot)*AdultT(Hot) | -0.027 | 0.043 | 574.199 | -0.627 | 0.531 |
|  | **Random effect** | **Variance** | **SD** |  |  |  |
|  | Family | 0.002 | 0.050 |  |  |  |
| **Main effects  model** | **Fixed effects** | **Estimate** | **SE** | **DF** | **t** | **P** |
|  | (Intercept) | 2.063 | 0.088 | 543.942 | 23.324 | <0.001 |
|  | DevT (Hot) | 0.059 | 0.028 | 627.055 | 2.096 | **0.036** |
|  | AdultT(Hot) | -0.815 | 0.022 | 574.440 | -37.734 | **<0.001** |
|  | Emergence weight | 0.083 | 0.016 | 593.952 | 5.305 | <0.001 |
|  | Block2 | 0.005 | 0.032 | 80.868 | 0.170 | 0.865 |
|  | Block3 | 0.113 | 0.036 | 123.959 | 3.121 | 0.002 |
|  | **Random effect** | **Variance** | **SD** |  |  |  |
|  | Family | 0.002 | 0.050 |  |  |  |

**Table S8.** The effects of developmental and adult temperature on **female fertility** (N=635). Modelled using a GLMM (Logit link function). “Full model” shows the parameter estimates and significance values for the model with two-way interactions (highlighted in grey), while the main-effects model shows parameter estimates and significance values for interpretation of only the main-effects (highlighted in grey).

| **Full model** | **Fixed effects** | **Estimate** | **SE** | **z** | **P** |
| --- | --- | --- | --- | --- | --- |
|  | (Intercept) | 19.212 | 833.519 | 0.023 | 0.982 |
|  | DevT (Hot) | -18.073 | 833.519 | -0.022 | 0.983 |
|  | AdultT (Hot) | -14.980 | 833.519 | -0.018 | 0.986 |
|  | Block 2 | 1.822 | 0.413 | 4.413 | <0.001 |
|  | Block 3 | -0.417 | 0.395 | -1.057 | 0.290 |
|  | DevT (Hot)*AdultT (Hot) | 14.544 | 833.519 | 0.017 | 0.986 |
| **Main-effects model** | **Fixed effects** | **Estimate** | **SE** | **z** | **P** |
|  | (Intercept) | 5.158 | 0.779 | 6.621 | <0.001 |
|  | Block 2 | 1.825 | 0.413 | 4.413 | <0.001 |
|  | Block 3 | -0.421 | 0.395 | -1.066 | 0.287 |
|  | DevT (Hot) | -3.976 | 0.735 | -5.405 | **<0.001** |
|  | AdultT (Hot) | -0.511 | 0.328 | -1.557 | 0.119 |

**Table S9**: The effects of developmental and adult temperature on **female lifetime reproductive success (LRS**) after accounting for selective disappearance (N=635). Modelled using a GLMM. “Full model” shows the parameter estimates and significance values for the model with two-way interactions (highlighted in grey), while the main-effects model shows parameter estimates and significance values for interpretation of only the main-effects (highlighted in grey). Log_e (y) data transformation used.

| **Full model** | **Fixed effects** | **Estimate** | **SE** | **z** | **P** |
| --- | --- | --- | --- | --- | --- |
|  | (Intercept) | 5.794 | 0.205 | 28.29 | <0.001 |
|  | Adult lifespan | -0.152 | 0.014 | -10.922 | <0.001 |
|  | DevT (Hot) | -1.329 | 0.111 | -11.934 | <0.001 |
|  | AdultT (Hot) | -1.35 | 0.145 | -9.328 | <0.001 |
|  | Block 2 | 1.038 | 0.11 | 9.412 | <0.001 |
|  | Block 3 | 0.18 | 0.126 | 1.427 | 0.154 |
|  | DevT (Hot)*AdultT (Hot) | 0.08 | 0.151 | 0.531 | 0.595 |
|  | **Random effects** | **Variance** | **SD** |  |  |
|  | Observation-level | 0.83 | 0.911 |  |  |
|  | Family | 0.036 | 0.189 |  |  |
| **Main-effects model** | **Fixed effects** | **Estimate** | **SE** | **z** | **P** |
|  | (Intercept) | 5.775 | 0.202 | 28.639 | <0.001 |
|  | Adult Lifespan | -0.153 | 0.014 | -10.924 | **<0.001** |
|  | DevT (Hot) | -1.288 | 0.081 | -15.981 | **<0.001** |
|  | AdultT (Hot) | -1.313 | 0.126 | -10.388 | **<0.001** |
|  | Block 2 | 1.038 | 0.11 | 9.413 | **<0.001** |
|  | Block 3 | 0.179 | 0.126 | 1.421 | **0.155** |
|  | **Random effects** | **Variance** | **SD** |  |  |
|  | Observation-level | 0.831 | 0.912 |  |  |
|  | Family | 0.035 | 0.189 |  |  |

**Table S10.** Test for differences in age independent traits between developmental temperatures within each Block. Significance determined using Tukey’s pairwise comparisons. Estimated marginal means may be on transformed data (See Table 1 for more information on transformations).

|  |  |  | | **Estimated marginal means (SE)** | |  |
| --- | --- | --- | --- | --- | --- | --- |
| **Trait** | **Sex** | **Block** | | **Ancestral** | **Hot** | **P** |
| Emergence success | Both | 1 | | 1.019 (0.168) | -0.503 (0.158) | **<0.001** |
|  |  | 2 | | 1.903 (0.187) | 1.406 (0.132) | **0.011** |
|  |  | 3 | | 1.122 (0.159) | -1.266 (0.130) | **<0.001** |
| Development time | Male | 1 | | 2.863 (0.007) | 2.623 (0.009) | **<0.001** |
|  |  | 2 | | 2.867 (0.006) | 2.530 (0.005) | **<0.001** |
|  |  | 3 | | 2.896 (0.006) | 2.632 (0.007) | **<0.001** |
|  | Female | 1 | | 5.650 (0.022) | 4.810 (0.028) | **<0.001** |
|  |  | 2 | | 5.610 (0.019) | 4.520 (0.016) | **<0.001** |
|  |  | 3 | | 5.730 (0.022) | 4.790 (0.026) | **<0.001** |
| Emergence weight | Male | | 1 | 3.380 (0.068) | 3.210 (0.088) | **0.076** |
|  |  | | 2 | 3.740 (0.058) | 3.570 (0.050) | **0.002** |
|  |  | | 3 | 4.000 (0.059) | 3.490 (0.070) | **<0.001** |
|  | Female | | 1 | 5.310 (0.082) | 4.40 (0.101) | **<0.001** |
|  |  | | 2 | 5.700 (0.071) | 4.86 (0.062) | **<0.001** |
|  |  | | 3 | 6.350 (0.081) | 4.47 (0.099) | **<0.001** |
| Adult lifespan | Male | | 1 | 14.400 (0.489) | 12.800 (0.652) | **0.027** |
|  |  | | 2 | 14.900 (0.345) | 16.200 (0.345) | **0.001** |
|  |  | | 3 | 14.700 (0.505) | 16.300 (0.505) | **0.007** |
|  | Female | | 1 | 2.030 (0.028) | 2.300 (0.040) | **<0.001** |
|  |  | | 2 | 2.180 (0.025) | 2.100 (0.021) | **0.018** |
|  |  | | 3 | 2.100 (0.034) | 2.400 (0.038) | **<0.001** |
| Fertility | Female | | 1 | 4.903 (0.749) | 0.927 (0.300) | **<0.001** |
|  |  | | 2 | 6.728 (0.783) | 2.752 (0.289) | **<0.001** |
|  |  | | 3 | 4.482 (0.732) | 0.505 (0.268) | **<0.001** |
| LRS | Female | | 1 | 3.900 (0.096) | 1.980 (0.137) | **<0.001** |
|  |  | | 2 | 4.240 (0.078) | 3.730 (0.066) | **<0.001** |
|  |  | | 3 | 4.250 (0.093) | 1.670 (0.137) | **<0.001** |

**Table S11:** The effects of developmental temperature, adult temperature, and age on **age-dependent** (**daily) fecundity of females** (N = 619). Modelled using GLMM (Log link function). “Full model” shows the parameter estimates and significance values for the model with two-way interactions (highlighted in grey), while the main-effects model shows parameter estimates and significance values for interpretation of only the main-effects (highlighted in grey). Referred to as “Model 1”.

| **Model** | **Fixed effects** | **Estimate** | **SE** | **z** | **P** |
| --- | --- | --- | --- | --- | --- |
| **Full** | (Intercept) | 3.928 | 0.199 | 19.696 | <0.001 |
|  | DevT(Hot) | -0.499 | 0.137 | -3.651 | <0.001 |
|  | AdultT(Hot) | 0.256 | 0.180 | 1.424 | 0.154 |
|  | Age | -0.135 | 0.023 | -5.789 | <0.001 |
|  | Age^2 | -0.008 | 0.002 | -4.592 | <0.001 |
|  | Adult Lifespan | -0.120 | 0.013 | -9.503 | <0.001 |
|  | Block2 | 1.029 | 0.110 | 9.327 | <0.001 |
|  | Block3 | 0.191 | 0.124 | 1.543 | 0.123 |
|  | DevT(Hot)*AdultT(Hot) | -0.099 | 0.229 | -0.434 | 0.664 |
|  | DevT(Hot)* Age | -0.310 | 0.031 | -10.037 | <0.001 |
|  | AdultT(Hot)* Age | -0.509 | 0.068 | -7.530 | <0.001 |
|  | DevT(Hot)*Age^2 | 0.019 | 0.002 | 9.142 | <0.001 |
|  | AdultT(Hot)*Age^2 | 0.025 | 0.008 | 3.065 | 0.002 |
|  | DevT(Hot)*AdultT(Hot)*Age | -0.065 | 0.102 | -0.635 | 0.525 |
|  | DevT(Hot)*AdultT(Hot)*Age^2 | 0.015 | 0.013 | 1.210 | 0.226 |
|  | **Random effects** | **Variance** | **SD** |  |  |
|  | ID | 0.594 | 0.771 |  |  |
|  | Family | 0.048 | 0.219 |  |  |
|  | **Fixed effects** | **Estimate** | **SE** | **z** | **P** |
| **Two-way** | (Intercept) | 3.977 | 0.195 | 20.426 | <0.001 |
|  | AdultT(Hot) | 0.206 | 0.146 | 1.413 | 0.158 |
|  | Age | -0.143 | 0.022 | -6.498 | <0.001 |
|  | DevT(Hot) | -0.598 | 0.103 | -5.781 | <0.001 |
|  | Age^2 | -0.007 | 0.002 | -4.565 | <0.001 |
|  | Adult Lifespan | -0.119 | 0.013 | -9.478 | <0.001 |
|  | Block2 | 1.024 | 0.110 | 9.273 | <0.001 |
|  | Block3 | 0.189 | 0.124 | 1.52 | 0.129 |
|  | AdultT(Hot)*Age | -0.533 | 0.051 | -10.451 | **<0.001** |
|  | DevT(Hot)*Age | -0.295 | 0.027 | -10.845 | **<0.001** |
|  | DevT(Hot)*Age^2 | 0.018 | 0.002 | 9.58 | **<0.001** |
|  | AdultT(Hot)*Age^2 | 0.031 | 0.006 | 4.893 | **<0.001** |
|  | **Random effects** | **Variance** | **SD** |  |  |
|  | ID | 0.593 | 0.77 |  |  |
|  | Family | 0.048 | 0.22 |  |  |

**Table S12:** Effect sizes (Hedge’s g) for eggs laid per day, when comparing females from hot adult temperatures (xH) versus ancestral adult (xA) temperatures. Positive sign indicates that females from hot adult temperature laid more eggs than females from ancestral adult temperatures, while negative sign indicates that females from ancestral adult temperatures laid more eggs. Hedge’s g interpretation: 0.2= small effect, 0.5 =medium effect, 0.8= large effect (Nakagawa & Cuthill, 2007).

| Age | Hedge's g |
| --- | --- |
| 1 | 0.819 |
| 2 | 0.496 |
| 3 | 0.134 |
| 4 | -0.478 |
| 5 | -0.732 |
| 6 | -0.722 |
| 7 | -0.724 |
| 8 | -0.639 |
| 9 | -0.456 |
| 10 | -0.123 |
| 11 | -0.469 |
| 12 | -0.406 |

**Table S13:** Comparison of models with different random effects structures, looking at the effects of developmental temperature adult temperature and age on daily fecundity of females. Comparisons are of the original model (i.e. model specified in Table S11) which allowed only for family and individual intercepts to vary, against a model which additionally allowed age dependent slopes to vary between families, against one which allowed age dependent slopes to vary between individuals, against one which allowed age-dependent slopes to vary between both families, and individuals . Models were compared using log-likelihood values following a Chi sq. distribution. Each model compared to the model above it. i.e. the next best fitting-model. The last model, i.e. the model which allows both between individual, and between family slopes to vary, provides the best fit to the data.

| **Model** | **Random effects** | **Df** | **AIC** | **BIC** | **logLik** | **deviance** | **Chisq** | Δ**DF** | **P** |
| --- | --- | --- | --- | --- | --- | --- | --- | --- | --- |
| Intercepts  allowed to vary between individuals  and families (Model 1) | 1\|ID + 1\|Family | 18 | 27336 | 27456 | -13650 | 27300 |  |  |  |
| Age dependent  slopes vary  between families | 1\|ID + Age\|Family | 20 | 27198 | 27332 | -13579 | 27158 | 141.68 | 2 | **<0.001** |
| Age dependent  slopes vary  between individuals | 1\| Family + Age\|ID | 20 | 26930 | 27064 | -13445 | 26890 | 267.901 | 0 | **<0.001** |
| Age dependent slopes vary between individuals and between families | Age\|ID + Age\|Family | 22 | 26924 | 27071 | -13440 | 26880 | 10.637 | 2 | **0.005** |

The model output of the analysis which allows slopes of both, different families and different females to vary, is presented below. This model is identical to the model presented in Table S11, except for its random effect structure.

**Table S14:** The effects of developmental temperature, adult temperature, and age on **male weight (mg)**. Modelled using LMMs. Three-way interactions (in grey). Full model uses males from both hot and ancestral adult temperatures (N=673).

| **Model** | **Fixed effects** | **Estimate** | **SE** | **DF** | **t** | **P** |
| --- | --- | --- | --- | --- | --- | --- |
| **Full** | (Intercept) | -0.038 | 0.042 | 983.40 | -0.896 | 0.370 |
|  | Block2 | 0.177 | 0.019 | 94.26 | 9.551 | <0.001 |
|  | Block3 | 0.247 | 0.019 | 109.20 | 12.707 | <0.001 |
|  | Adult lifespan | 0.020 | 0.001 | 740.00 | 16.280 | <0.001 |
|  | Age | -0.056 | 0.003 | 5097.00 | -19.385 | <0.001 |
|  | Emergence.weight | 0.837 | 0.011 | 1055.00 | 78.470 | <0.001 |
|  | DevT(Hot) | -0.041 | 0.017 | 1357.00 | -2.445 | 0.015 |
|  | AdultT(Hot) | 0.297 | 0.023 | 1335.00 | 12.976 | <0.001 |
|  | Age^2 | 0.002 | 0.000 | 4726.00 | 34.532 | <0.001 |
|  | Age*Emergence.weight | -0.017 | 0.001 | 5187.00 | -26.469 | <0.001 |
|  | DevT(Hot)*AdultT(Hot) | -0.013 | 0.024 | 1991.00 | -0.528 | 0.598 |
|  | DevT(Hot)*Age | 0.017 | 0.002 | 4707.00 | 8.333 | <0.001 |
|  | AdultT(Hot)*Age | -0.205 | 0.005 | 4767.00 | -41.096 | <0.001 |
|  | DevT(Hot)*Age^2 | -0.001 | 0.000 | 4757.00 | -6.867 | <0.001 |
|  | AdultT(Hot)*Age^2 | 0.011 | 0.001 | 4900.00 | 20.910 | <0.001 |
|  | DevT(Hot)*AdultT(Hot)*Age | 0.036 | 0.007 | 4745.00 | 5.100 | **<0.001** |
|  | DevT(Hot)*AdultT(Hot)*Age^2 | -0.002 | 0.001 | 4872.00 | -2.831 | **0.005** |
|  | **Random effect** | **Variance** | **SD** |  |  |  |
|  | ID | 0.011 | 0.107 |  |  |  |
|  | Family | 0.002 | 0.044 |  |  |  |

**Table S15:** Post-hoc test conducted on age-dependent male weight, shown in model S14. The effects of developmental temperature on age-dependent male weight are analysed for males who only experience hot adult temperatures, to compare AH (*Ancestral Developmental and Hot Adult*) and HH (*Hot Developmental and Hot Adult*) treatments. (N=354)

| Fixed effects | Estimate | SE | DF | t | P |
| --- | --- | --- | --- | --- | --- |
| (Intercept) | -0.117 | 0.054 | 847.900 | -2.189 | 0.029 |
| Age | -0.180 | 0.010 | 1468.000 | -17.501 | <0.001 |
| Emergence weight | 0.883 | 0.015 | 830.800 | 60.551 | <0.001 |
| DevT (Hot) | -0.054 | 0.017 | 999.200 | -3.119 | 0.002 |
| Age^2 | 0.013 | 0.001 | 1425.000 | 24.618 | <0.001 |
| Block 2 | 0.153 | 0.020 | 83.070 | 7.678 | <0.001 |
| Block 3 | 0.206 | 0.021 | 92.310 | 9.934 | <0.001 |
| Adult lifespan | 0.047 | 0.003 | 347.900 | 15.869 | <0.001 |
| Age*Emergence weight | -0.039 | 0.002 | 1463.000 | -15.708 | <0.001 |
| DevT (Hot)*Age | 0.049 | 0.007 | 1368.000 | 7.011 | **<0.001** |
| DevT (Hot)*Age^2 | -0.003 | 0.001 | 1447.000 | -3.596 | **<0.001** |
| Random effects | Variance | SD |  |  |  |
| ID | 0.005 | 0.071 |  |  |  |
| Family | 0.001 | 0.039 |  |  |  |

**Table S16:** Post-hoc test conducted on age-dependent male weight, shown in model S14. The effects of developmental temperature on age-dependent male weight are analysed for males who only experience ancestral adult temperatures, to compare AA (*Ancestral Developmental and Ancestral Adult*) and HA (*Hot Developmental and Ancestral Adult)* treatments. (N=319)

| Fixed effects | Estimate | SE | DF | t | P |
| --- | --- | --- | --- | --- | --- |
| (Intercept) | 0.053 | 0.057 | 459.606 | 0.918 | 0.359 |
| Age | -0.062 | 0.003 | 3435.744 | -21.399 | <0.001 |
| Emergence weight | 0.834 | 0.016 | 442.730 | 51.859 | <0.001 |
| DevT (Hot) | -0.028 | 0.018 | 560.570 | -1.558 | 0.120 |
| Age^2 | 0.002 | 0.000 | 3338.217 | 35.750 | <0.001 |
| Block 2 | 0.165 | 0.026 | 91.999 | 6.384 | <0.001 |
| Block 3 | 0.264 | 0.027 | 105.572 | 9.736 | <0.001 |
| Adult lifespan | 0.016 | 0.001 | 352.191 | 11.167 | <0.001 |
| Age*Emergence weight | -0.016 | 0.001 | 3451.102 | -23.115 | <0.001 |
| DevT (Hot)*Age | 0.018 | 0.002 | 3329.344 | 9.033 | **<0.001** |
| DevT (Hot)*Age^2 | -0.001 | 0.000 | 3357.817 | -7.442 | **<0.001** |
| Random effects | Variance | SD |  |  |  |
| ID | 0.013 | 0.113 |  |  |  |
| Family | 0.004 | 0.06 |  |  |  |

**Table S17:** The effects of developmental temperature and adult temperature on **male age-dependent mortality** (N = 689). Modelled using a Cox- proportional hazards mixed effects model. “Full model” shows the parameter estimates and significance values for the model with two-way interactions (highlighted in grey), while the main-effects model shows parameter estimates and significance values for interpretation of only the main-effects (highlighted in grey).

| **Model** | **Fixed effects** | **Estimate** | **z** | **P** |
| --- | --- | --- | --- | --- |
| Full | DevT (Hot) | -0.153 | -1.260 | 0.210 |
|  | AdultT (Hot) | 3.161 | 19.110 | <0.001 |
|  | Block2 | -0.094 | -0.730 | 0.460 |
|  | Block3 | -0.474 | -3.240 | 0.001 |
|  | DevT (Hot) * AdultT (Hot) | 0.090 | 0.530 | 0.590 |
|  | **Random effect** | **SD** | **Variance** |  |
|  | Family | 0.296 | 0.087 |  |
| **Model** | **Fixed effects** | **Estimate** | **z** | **P** |
| Main effects | DevT (Hot) | -0.107 | -1.260 | 0.210 |
|  | AdultT (Hot) | 3.207 | 22.540 | **<0.001** |
|  | Block2 | -0.094 | -0.730 | 0.460 |
|  | Block3 | -0.476 | -3.250 | 0.001 |
|  | **Random effect** | **SD** | **Variance** |  |
|  | Family | 0.297 | 0.088 |  |

**Table S18:** The effects of developmental temperature and adult temperature on **female age-dependent mortality** (N = 635). Modelled using a Cox- proportional hazards mixed effects model. “Full model” shows the parameter estimates and significance values for the model with two-way interactions (highlighted in grey), while the main-effects model shows parameter estimates and significance values for interpretation of only the main-effects (highlighted in grey).

| **Model** | **Fixed effects** | **Estimate** | **z** | **P** |
| --- | --- | --- | --- | --- |
| Full | DevT (Hot) | -0.656 | -0.550 | 0.590 |
|  | AdultT (Hot) | 4.302 | 19.610 | <0.001 |
|  | Block2 | -0.878 | -5.690 | <0.001 |
|  | Block3 | -0.793 | -5.060 | <0.001 |
|  | DevT (Hot) * AdultT (Hot) | 0.035 | 0.220 | 0.820 |
|  | **Random effect** | **SD** | **Variance** |  |
|  | Family | 0.367 | 0.135 |  |
| **Model** | **Fixed effects** | **Estimate** | **z** | **P** |
| Main-effects | DevT (Hot) | -0.047 | -0.540 | 0.590 |
|  | AdultT (Hot) | 4.319 | 21.150 | **<0.001** |
|  | Block2 | -0.879 | -5.700 | <0.001 |
|  | Block3 | -0.796 | -5.080 | <0.001 |
|  | **Random effect** | **SD** | **Variance** |  |
|  | Family | 0.366 | 0.134 |  |

D. **Note on Selective disappearance of beetles during emergence**

Out of the three experimental blocks, the hot treatments in blocks 1 and 3 were set up at 36 ˚C while the hot treatments in block 2 were set up at 33 ˚C. This difference was the result of a technical failure in one of our incubators, which meant that beetles in block 2 were housed in a controlled temperature room whose temperature could not exceed 33 ˚C.

One consequence of this temperature difference between blocks was that the developmental environment in block 2 was less stressful for the beetles than blocks 1 and 3. 94 out of 260 eggs emerged as adult beetles from the hot developmental treatment in block 1 (36%), 460 out of 600 eggs emerged as adult beetles from the hot developmental treatment in block 2 (77%), and 134 out of 600 eggs emerged as adult beetles from the hot developmental treatment in block 3 (22%) (Table S1).

We also found that hot developmental temperatures increased adult lifespan (after accounting for the effects of weight) (Table S4). This result is somewhat counterintuitive given hot temperatures are expected to be stressful. To investigate whether this could be due to non-random death of beetle larvae and pupae during development in the hot treatment, leading to selective disappearance of poor condition individuals we compared the egg to adult emergence success and lifespan of individuals between the three blocks. Because emergence success in the hot developmental temperature was greater in block 2 than it was in blocks 1 and 3, we expected that if selective disappearance during development in hot temperatures was occurring, its magnitude should be greater in blocks 1 and 3 than in block 2. Consequently, if selective disappearance during development caused mean lifespan of adult beetles from hot developmental temperatures to increase, we should have expected hot developmental temperatures in blocks 1 and 3 to have a similar (positive) effect on adult lifespan of beetles and in block 2 to have a different effect (negative). But this is not we found (Table S4).

Instead, for males, we found that hot developmental temperatures in both blocks 2 and 3 had a significantly positive effect on lifespan while hot temperatures in block 1 had a significantly negative effect on lifespan. Furthermore, the effects of block 2 on adult lifespan was more similar to the effects of block 3 than the effects of block 3 were to that of block 1. This is not what we would have expected if hotter temperatures were leading to selective disappearance of beetles at the larval stage (Table S10).

On the other hand, for females, we indeed found that hot developmental temperatures in blocks 1 and 3 (which were at a hotter temperature than block 2 i.e. 36 ˚C vs 33 ˚C) increased female lifespan compared to block 2. This could mean that it is due to selective disappearance of lower quality females at the larval stage, that female lifespan is positively affected by hot developmental temperatures.

We thus suggest that while for males, it is unlikely that selective disappearance could have caused an increase in adult lifespan under hot developmental temperatures, the opposite may be true for females.

E. **Note on selective disappearance of lighter males during adulthood**

To further examine whether the effects of an increase in average weight seen towards the end of life in males was due to selective disappearance of lighter males, we visualised individual-level changes in weight with age. First, we binned males into categories based on their lifespans to plot their age-dependent weight as smoothed splines (Figure S1). This showed a strong correlation of males of a lower emergence weight having shorter lifespans.

Figures S1 shows that there is a decline in weight with age for each curve in each treatment, and that heavier individuals live longer, thus any increase in weight with advancing age, as seen in Figure 4, and due to a quadratic function of age in Table S14, can be attributed to selective disappearance of lighter beetles rather than an actual increase in weight of individuals with age.

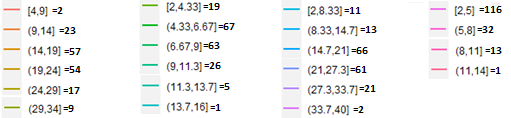

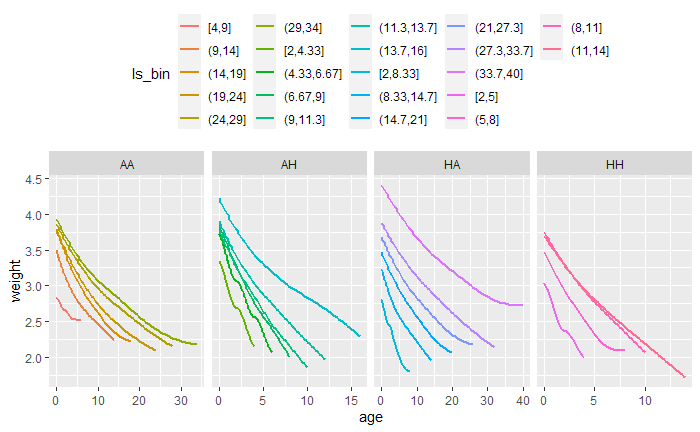

Figure S1. Change in weight (mg) with adult age (in days), of males, with plot binned by male Adult lifespan ((age range for each bin] = sample size of males in each bin). *Ancestral Developmental and Ancestral Adult (****AA****), Ancestral Developmental and Hot Adult (****AH****), Hot Developmental and Ancestral Adult (****HA****)*, *and Hot Developmental and Hot Adult (****HH****)* temperatures*.* Each smoothed spline is created using the average weight of males in that given lifespan group. Males which have higher weights at adult age 0 (emergence) live longer than males which have lower emergence weights. 4 to 6 bins were created for each treatment because these allowed the clearest interpretation of curves visually, with the least amount of lines crossing over.
